# Oxytocin increases the pleasantness of affective touch and orbitofrontal cortex activity independent of valence

**DOI:** 10.1101/2020.02.26.965830

**Authors:** Yuanshu Chen, Benjamin Becker, Yingying Zhang, Han Cui, Jun Du, Jennifer Wernicke, Keith M. Kendrick, Shuxia Yao

**Affiliations:** The Clinical Hospital of Chengdu Brain Science Institute, MOE Key Laboratory for NeuroInformation, Center for Information in Medicine, University of Electronic Science and Technology of China, Chengdu, Sichuan 611731, China; Department of Molecular Psychology, Institute of Psychology and Education, Ulm University, Ulm, Germany

**Keywords:** affective touch, oxytocin, valence, orbitofrontal cortex

## Abstract

Touch plays a crucial role in affiliative behavior and social communication. The neuropeptide oxytocin is released in response to touch and may act to facilitate the rewarding effects of social touch. However, no studies to date have determined whether oxytocin facilitates behavioral or neural responses to non-socially administered affective touch and possible differential effects of touch valence. In a functional MRI experiment using a randomized placebo-controlled, within-subject design in 40 male subjects we investigated the effects of intranasal oxytocin (24IU) on behavioral and neural responses to positive, neutral and negative valence touch administered to the arm via different types of materials at a frequency aimed to optimally stimulate C-fibers. Results showed that oxytocin significantly increased both the perceived pleasantness of touch and activation of the orbitofrontal cortex independent of touch valence. The effects of OT on touch-evoked orbitofrontal activation were also positively associated with basal oxytocin concentrations in blood. Additionally, anterior insula activity and the functional connectivity between the amygdala and right anterior insula were enhanced only in response to negative valence touch. Overall, the present study provides the first evidence that oxytocin may facilitate the rewarding effects of all types of touch, irrespective of valence.

## 1. Introduction

Touch, including both the discriminative and affective touch, is one of the dominant sensory modalities for human beings to perceive the external world and is thus of crucial importance in human development (Field, 2014; Schirmer and Adolphs, 2017). While discriminative touch serves to detect and identify physical aspects such as pressure, affective touch conveys more qualitative emotional aspects. Affective touch in the context of social interactions forms the basis of social affiliation and shapes social reward and emotional regulation throughout an individual’s life (Cascio et al., 2019; McGlone et al., 2014; Morrison et al., 2010). Recent studies have emphasized the importance of slow unmyelinated low-threshold C touch (CT) fibers in processing pleasant affective touch (Ackerley et al., 2014a; Croy et al., 2016; Pawling et al., 2017). These particular fibers mainly respond to gentle, slow, caress-like stroking targeting hairy skin (Ackerley et al., 2014a, 2014b; Löken et al., 2009; Morrison et al., 2010). Previous studies have shown that CT-targeted touch mainly influences reponses in regions of the ‘social brain’ such as the amygdala, insula, orbitofrontal cortex (OFC) and superior temporal sulcus (STS) (Björnsdotter et al., 2014, 2010; Croy et al., 2016; Gordon et al., 2013b; Mcglone et al., 2012; Morrison et al., 2011).

Oxytocin (OT), a hypothalamic neuropeptide, is a key modulator in regulating social behavior and affective processing across species (Bartz et al., 2011; Feldman, 2012; Kendrick et al., 2017; Olff et al., 2013). Previous studies have also demonstrated a close association between OT and CT-targeted pleasant social touch (Walker et al., 2017). More specifically, it has been reported that stroking touch and massage facilitate endogenous OT release in both animals (Agren et al., 1995; Lund et al., 2002; Uvnäs-Moberg et al., 2015) and humans (Holt-Lunstad et al., 2008; Li et al., 2019; Light et al., 2005; Morhenn et al., 2012) and activates oxytocin neurons in the paraventricular nucleus (Okabe et al., 2015). Moreover, one recent fMRI study has reported that intranasal OT increases the perceived pleasantness of positive social touch applied to the leg as well as insula and OFC activation (Scheele et al., 2014). This later study in male subjects only found an effect of positive social touch when subjects thought the touch was administered by a female and not by a male, even though unbeknown to them the touch was always administered by the same female. Thus, the effects of OT on increasing the pleasantness of touch may be influenced by psychological factors affecting perceived valence. However, no studies have directly investigated the importance of either touch valence on the influence of OT on behavioral and neural responses or whether it can also modulate responses to touch administered in a non-social context.

While CT-fibers seem primarily to respond to pleasant touch (Croy et al., 2016), there is also some evidence of OT’s analgesic effects that OT is released following painful/aversive responses to stimuli applied to the skin and that perceived unpleasantness of stimuli can be reduced by intranasal OT (see Boll et al., 2018; Walker et al., 2017). Indeed, there is considerable overlap between brain regions activated by pain and touch (Lui et al., 2008). It is therefore possible that OT can both enhance the perceived pleasantness of pleasant touch and reduce the aversiveness of unpleasant touch. The current study has therefore investigated whether OT can enhance the pleasantness of both pleasant and aversive touch by influencing brain reward mechanisms and other regions responsive to affective touch.

The present study employed a CT-targeted affective touch paradigm combined with the fMRI to investigate whether OT modulates behavioral and neural responses to different valences of affective touch administered in a non-social context using different materials. Touch applied using such materials can also activate CT-fibers and brain regions associated with social touch if administered at the optimum slow velocity (Voos et al., 2013) and offers the advantage of being able to use different materials to influence the perceived valence without including a social intention factor (Kirsch et al., 2018; von Mohr et al., 2018). Thus, we used this CT-targeted affective touch strategy to achieve better control over possible confounding effects of perceived social intention. In addition, we investigated whether the effects of OT are modulated by autistic traits and attitudes to positive social touch given findings from a previous study reporting a negative association (Scheele et al., 2014) and that autistic individuals can exhibit aberrant behavioral or neural responses to both positive and negative valence affective touch (Cascio et al., 2012; Green et al., 2015; Kaiser et al., 2016). Some studies have reported associations between autistic symptom severity and basal concentrations of OT in blood in both healthy (Li et al., 2019) and clinical populations (Zhang et al., 2016), and that effects of intranasal OT in autistic individuals can also be associated with them (Gordon et al., 2013a; Parker et al., 2017). We therefore also investigated potential correlations between basal blood OT concentrations and the behavioral and neural effects of intranasal OT on responses to affective touch.

We hypothesized that if OT can both increase the pleasantness of positive valence touch and reduce aversion for unpleasant touch then it would have similar behavioral and brain reward system effects independent of valence. We additionally hypothesized that OT might alter responsiveness to negative valence touch by influencing activity in regions such as the insula which respond more selectively to aversive stimuli (see Gogolla, 2017). Finally, we hypothesized that both behavioral and neural effects of OT on affective touch would be modulated by trait autism and sensitivity to social touch and positively associated with basal OT concentrations in blood.

## 2. Methods and Materials

### 2.1 Participants

A total of 50 healthy male subjects (mean age = 21.19 years old, sd = 2.62) were recruited by local advertisement to participate in the present study. All subjects had normal or corrected-to-normal vision and were free of current or a history of psychiatric or physical disorders. Ten subjects were excluded due to either failure to attend the second part of the experiment (5 subjects) or excessive head movement in either one or both of the scanning sessions (5 subjects). Written informed consent was obtained from all subjects before the experiment. All procedures were in accordance with the latest revision of the declaration of Helsinki and approved by the local ethics committee at the University of Electronic Science and Technology of China.

### 2.2 Procedures

A randomized, double-blind, placebo-controlled within-subject pharmaco-fMRI design was used in the present study. Subjects underwent two fMRI assessments and were randomly administered either OT (24 IU; Oxytocin Spray, Sichuan Meike Pharmaceutical Co. Ltd, China) or placebo (PLC, identical ingredients other than OT) intranasal spray 45 min before the experiment following a standardized protocol (Guastella et al., 2013). Immediately prior to intranasal administration blood samples (6 ml) were collected from all subjects by venipuncture for measurement of basal OT concentrations. Treatment-order was counterbalanced across participants and the 2 visits were scheduled with an interval of at least 2 weeks. The debrief interview after each session showed that subjects were unable to guess better than chance whether they had received OT or PLC (χ^2^ = 0.77, P = 0.38). Prior to treatment and MRI scanning, subjects completed Chinese versions of validated questionnaires on individual personality traits and attitude towards personal touch. Personality trait measures included the Beck Depression Inventory II (BDI-II) (Beck et al., 1996), the State-Trait Anxiety Inventory (STAI) (Spielberger et al., 1983), the Autism Spectrum Quotient (AQ) (Baron-Cohen et al., 2001), the Liebowitz Social Anxiety Scale (LSAS) (Heimberg et al., 1999) and the Empathy Quotient (EQ) (Baron-Cohen and Wheelwright, 2004). Individual attitudes and sensitivity to touch were assessed using the Social Touch Questionnaire (STQ) (Wilhelm et al., 2001). Additionally, to control for possible confounding effects of treatment on mood across the experiment, subjects completed the Positive and Negative Affect Schedule (PANAS) (Watson et al., 1988) before and after treatment (pre/post-treatment) as well as after scanning (post-scan).

### 2.3 Affective touch task

In line with the pharmaco-dynamic profile of intranasal OT administration (Paloyelis et al., 2016; Spengler et al., 2017), the affective touch task started 45 min after treatment. During fMRI assessment, moderate-force touch stimulation was administered to an 8-cm marked zone on the left dorsal forearm by a trained experimenter at a consistent velocity of 8 cm/s, which has been shown to be the best rate to activate CT fibers (Löken et al., 2009; Olausson et al., 2010; Voos et al., 2013). To examine effects of OT on different valences of affective touch, three types of touch stimulation conditions were employed based on previous studies (Cascio et al., 2012; Essick et al., 2010): a plastic rough mesh material (negative valence) vs. a burlap (neutral) vs. a soft goat hair brush (positive valence) (Figure 1). Pleasantness and roughness of touch materials were rated by an independent sample of 12 subjects using a 9-point Likert scale (- 4 = extremely unpleasant or extremely rough, and + 4 = very pleasant or not rough at all). For pleasantness, mean ratings were – 1.63 ± 1.19 for negative, 0.20 ± 1.16 for neutral and + 2.46 ± 0.62 for positive conditions and for roughness, – 2.42 ± 0.82 for negative, – 1.08 ± 1.04 for neutral and + 1.50 ± 1.02 for positive conditions. Width and thickness of different touch materials were additionally controlled across conditions. The touch region on the forearm was marked before the experiment and the experimenter was instructed to vary the exact start and end point of each stroke by a few millimeters in order to minimize receptor fatigue (cf. Cascio et al., 2012).

**Figure 1.**
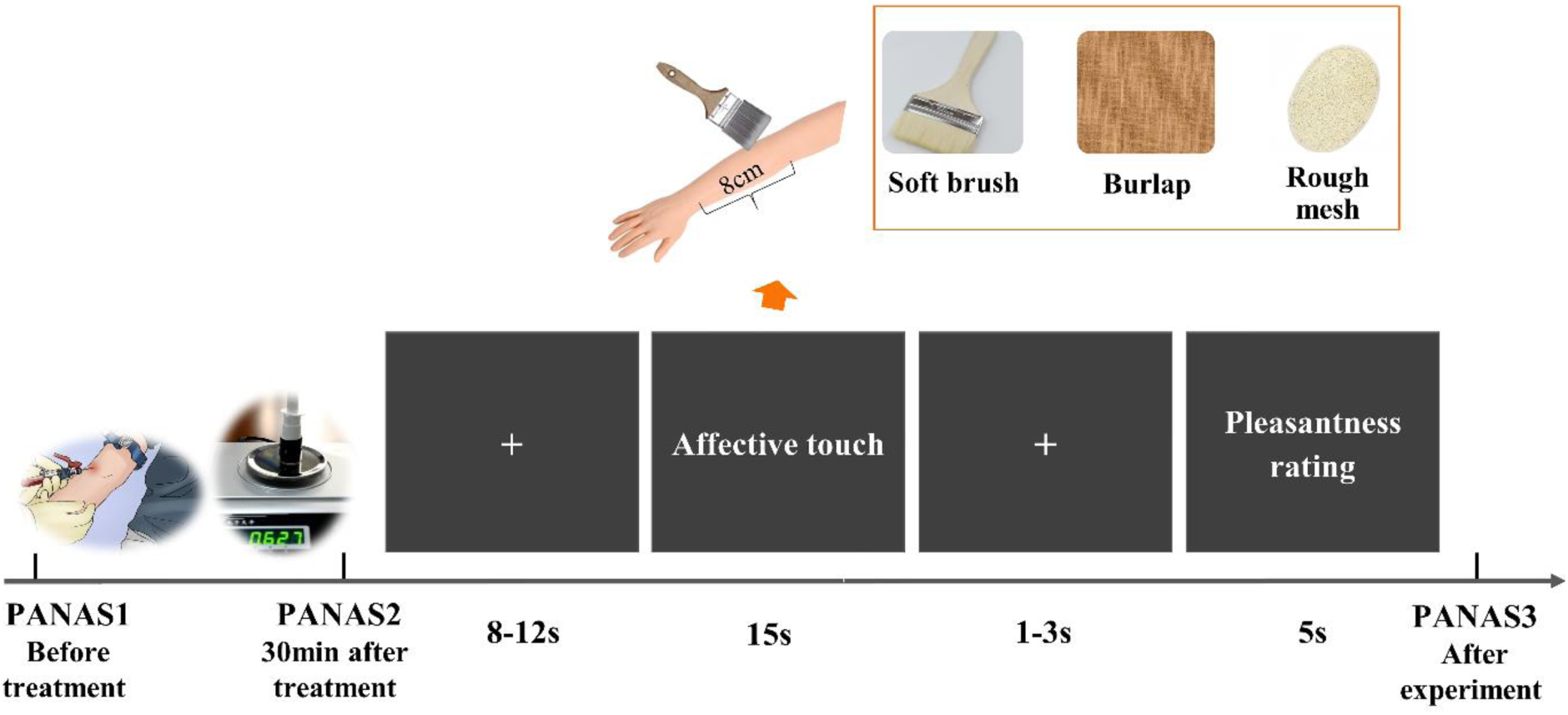
The fMRI affective touch task. Positive, neutral and negative valence touch stimuli used stroking via a soft brush, burlap and scratchy mesh respectively. Intranasal administration blood samples (6 ml) were collected for each subject before the treatment administration. Subjects completed the positive and negative affective schedule (PANAS) before and after intranasal oxytocin or placebo treatment and additionally after the task to measure mood.

There were 2 functional runs in the affective touch task. Each run consisted of 18 blocks with 6 blocks in each touch valence condition. In each 15-s block, participants received 15 tactile touch stimuli followed by a jittered interval of 1 – 3 s (mean 2s). Next participants were required to rate their perceived pleasantness of each applied touch block within 5 s on a 9-point Likert scale ranging from – 4 (extremely unpleasant) to + 4 (extremely pleasant). Different valence blocks were presented in a pseudorandom order with jittered fixation (mean 10 s). The stimulation periods and touch conditions were indicated to the experimenter by a second monitor. To ensure participants could not see the touch stimuli or the experimenter a blanket was used to cover the bore of the scanner. In total the experiment lasted 19.5 min (Figure 1).

### 2.4 Behavioral data analyses

The effects of treatment on behavioral pleasantness ratings were analyzed by repeated-measure analysis of variance (ANOVA) with treatment and valence as within-subject factors (using SPSS version 20). Partial eta-squared and Cohen’s d were calculated as measures of effect size. All reported P values were two tailed, and P < 0.05 was considered as significant.

### 2.5 Images acquisition and data analyses

Images were acquired using a 3.0 Tesla GE Discovery MR750 system (General Electric Medical System, Milwaukee, WI, USA). Time series of volumes were acquired using a T2*- weighted echo-planar pulse sequence (repetition time: 2000 ms; echo time: 30 ms; slices: 43; slice thickness: 3.2 mm; field of view: 220 × 220 mm; acquisition matrix: 64 × 64; flip angle: 90°). To control for anatomical abnormalities and increase normalization accuracy during preprocessing, high revolution T1-weighted anatomical images were acquired obliquely with a 3D spoiled gradient echo pulse sequence (repetition time: 6 ms; echo time: minimum; slices: 156; slice thickness: 1 mm; field of view: 256 × 256 mm; acquisition matrix: 256 × 256; flip angle: 12°).

Images were processed using SPM12 software (Wellcome Department of Cognitive Neurology, London; http://www.fil.ion.ucl.ac.uk/spm/spm12) (Friston et al., 1994). The first five functional volumes were discarded to achieve magnet-steady images. Realignment was then conducted on the remaining functional images to correct for head movement using the 6- parameter rigid body algorithm. After co-registering the mean functional image and the T1 image, the T1 image was segmented to determine the parameters for normalizing the functional images to the Montreal Neurological Institute (MNI) space. The normalized images were finally spatially smoothed using a Gaussian kernel (8 mm full-width at half maximum).

On the first level, 12 experimental conditions were modelled in the design matrix (negative, positive and neutral touch following OT and PLC treatment separately and pleasantness ratings for each of these touch conditions) and convolved with the canonical hemodynamic response function. The six head-motion parameters were additionally included as regressors in the design matrix to further control for head motion. Contrast images for each type of touch and additionally across all valences were created.

On the second level, a flexible factorial design was employed to examine the main effect of touch valence and treatment and the interaction. For whole brain analyses, a significance threshold of P < 0.05 false discovery rate (FDR) corrected at peak level was set for multiple comparisons. Furthermore, based on previous studies investigating neural mechanisms underlying affective tactile touch (Voos et al., 2013) and brain regions involved in the modulatory effects of intranasal OT on social touch (Scheele et al., 2014), we also performed a region of interest (ROI) analysis for regions including the medial and lateral amygdala and STS previously associated with social affective touch but which could not survive at whole brain level. These ROIs were anatomically defined based on the Automated Anatomic Labeling atlas (Tzourio-Mazoyer et al., 2002) or the Human Brainnetome Atlas (Fan et al., 2016). Within these a priori ROIs, a threshold of P < 0.05 family-wise error (FWE) peak-level correction was set for multiple comparisons using small volume correction (SVC). Brain-behavior correlations were calculated using Spearman correlation for both the OT and PLC conditions. Parameter estimates used for plotting and brain-behavior association analyses were extracted for each subject from a 6-mm sphere centered on the peak voxel within corresponding ROIs.

### 2.6 Oxytocin immunoassay

Blood collection and OT assays were in accordance with a previous study (Li et al., 2019). Briefly, venous blood was collected into 6 ml EDTA tubes and following centrifugation plasma stored at −80°C until OT analysis. Oxytocin concentrations in 1-ml plasma samples were measured in triplicate using a commercial ELISA assay (Cayman Chemical, Ann Arbor, Michigan USA Kit 500440) and a prior extraction step was performed in accordance with the manufacturers recommended protocol. All samples had detectable OT concentrations. The intra- and inter-assay coefficients of variation were 6 and 8% respectively. The manufacturer’s reported cross-reactivity of the antibody with related neuropeptides, such as vasopressin and vasotocin, is < 0.01%.

## 3. Results

### 3.1 Personality trait and mood questionnaires

Questionnaire scores on personality traits are reported in Table S1. Analyses on PANAS scores showed no significant influence of treatment *per se* but there was a significant increase in the positive mood scores (P = 0.04, Cohen’s d = 0.32) after the fMRI scanning relative to the post-treatment measurement in the OT group (see supplemental materials; Figure S1). This would seem to indicate that subjects under OT generally experienced a more positive mood following the affective touch paradigm than under PLC.

### 3.2 Effects of OT on pleasantness of affective touch

For the effects of OT on pleasantness ratings of touch, a repeated-measurement ANOVA with valence (negative vs. neutral vs. positive) and treatment (OT vs. PLC) as within-subject factors revealed a significant main effect of valence (F(2, 78) = 138.59, P < 0.001, η^2^ = 0.78). The post hoc Bonferonni corrected analysis showed that subjects rated the positive valence touch (2.12 ± 0.13) as more pleasant than both neutral (0.46 ± 0.14) and negative valence touch (−0.65 ± 0.15) (Ps < 0.001). Furthermore, there was a significant main effect of treatment (F(1, 39) = 5.77, P = 0.02, η^2^ = 0.13), with subjects following OT treatment rating all touch valences as more pleasant compared to the PLC treatment (M _OT_ = 0.75 ± 0.10, M _PLC_ = 0.53 ± 0.12). While there was no significant interaction between valence and treatment (P = 0.57) an exploratory post-hoc analysis showed that only the negative valence stimuli showed a significant increase in pleasantness ratings under OT (negative: P = 0.045, Cohen’s d = 0.30; positive and neutral: Ps > 0.11) (see Figure 2).

**Figure 2.**
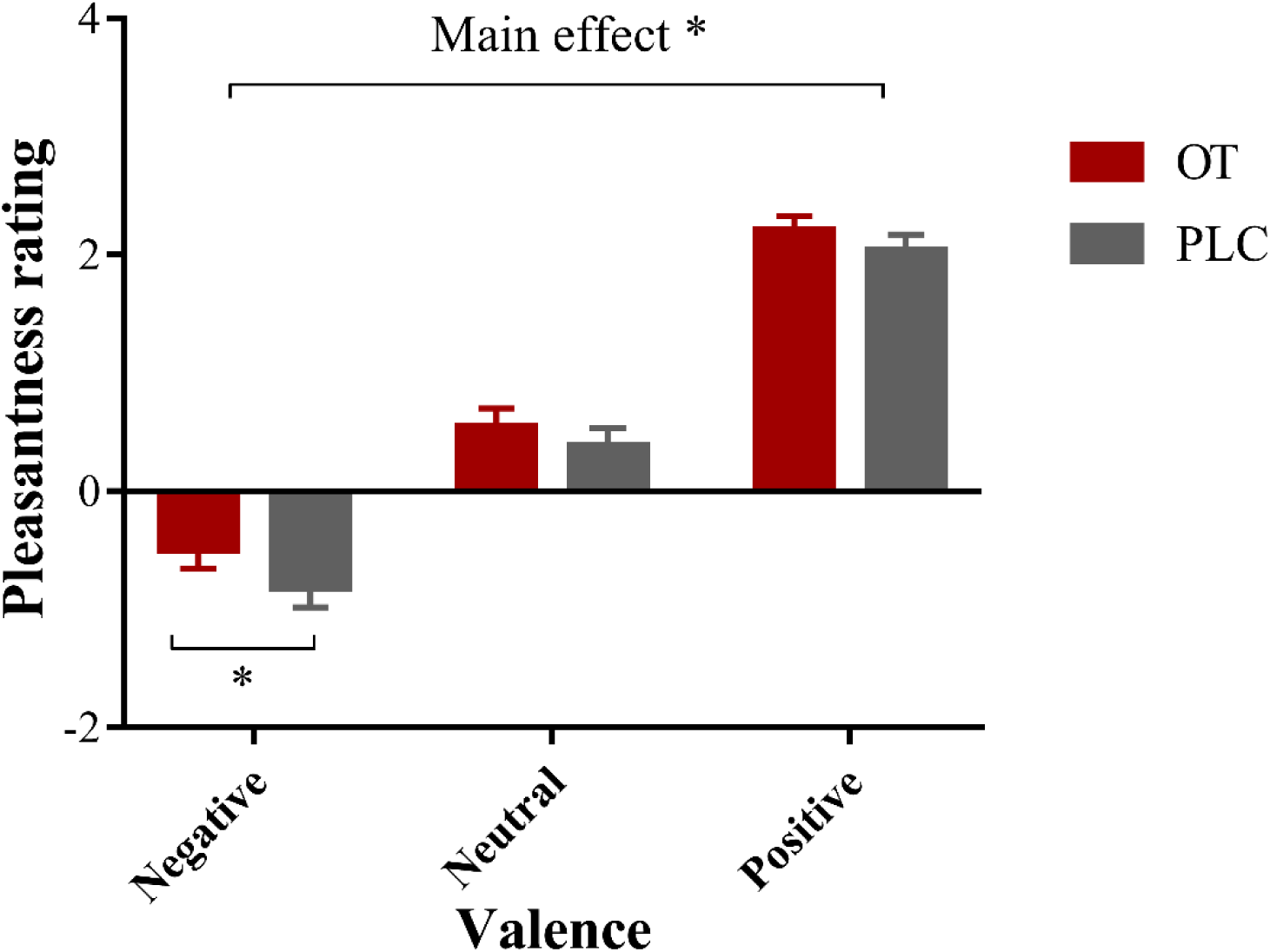
Pleasantness rating scores for affective touch stimuli under oxytocin (OT) and placebo (PLC) treatment. Histograms show mean ± SEM rating scores for the negative, neutral and positive valence touch stimuli. The line over the histograms indicates a significant main effect of treatment and within the negative valence condition also a significant effect of OT. Error bars show standard errors. ^*^P < 0.05.

### 3.3 Effects of OT on neural responses to affective touch

At whole brain level, increased activation was found in a number of different brain regions following positive relative to neutral valence touch, including the medial prefrontal cortex (mPFC), inferior frontal gyrus (IFG), inferior parietal lobe (IPL), anterior cingulate cortex (ACC), STS, insula, precentral gyrus and primary somatosensory cortex (SI) (P_FDR_ < 0.05). Positive valence touch also induced stronger activations in a number of brain regions including the OFC, mPFC, STS and insula compared to the negative valence touch (P_FDR_ < 0.05). A region of the precentral gyrus was also more activated by neutral than by positive or negative touch (P_FDR_ < 0.05). No regions exhibited greater response to negative compared to neutral or positive touch. Results further revealed a significant main effect of treatment across touch conditions in the OFC (P_FDR_ < 0.05; Figure 3a). A separate examination of OT’s effects on each touch valence revealed increased left AI activity (P_FDR_ < 0.05; Figure 3b) in response to negative touch following OT compared to PLC treatment, but not to the positive or neutral touch conditions. A summary of all of these whole brain findings is provided in Table 1. No other significant neural activation differences were observed at whole brain level with the same corrected threshold. There were also no significant effects of OT on neural responses in the a priori ROIs (STS and medial and lateral amygdala, P_FWE_ < 0.05, SVC).

**Table 1.**
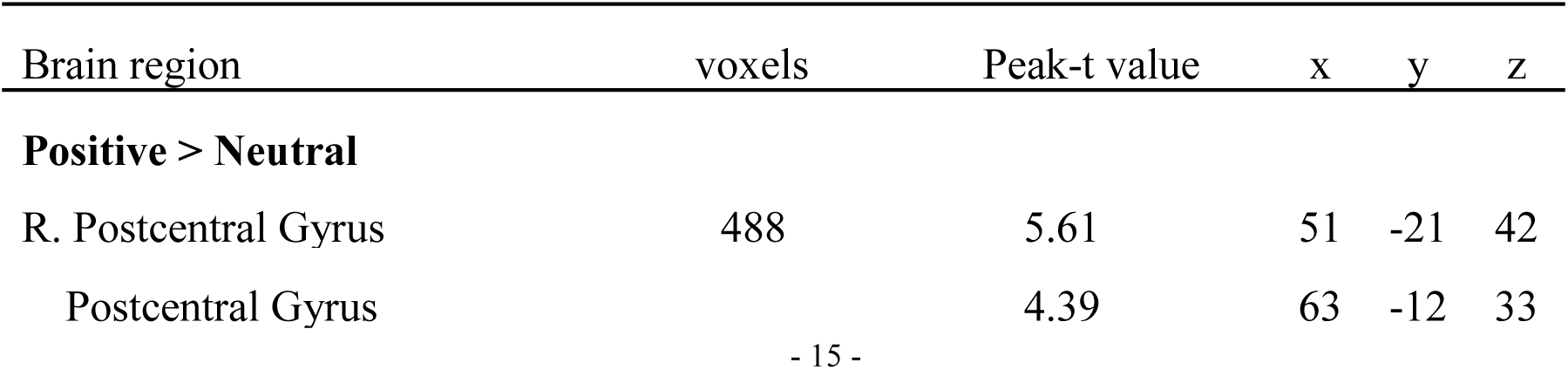
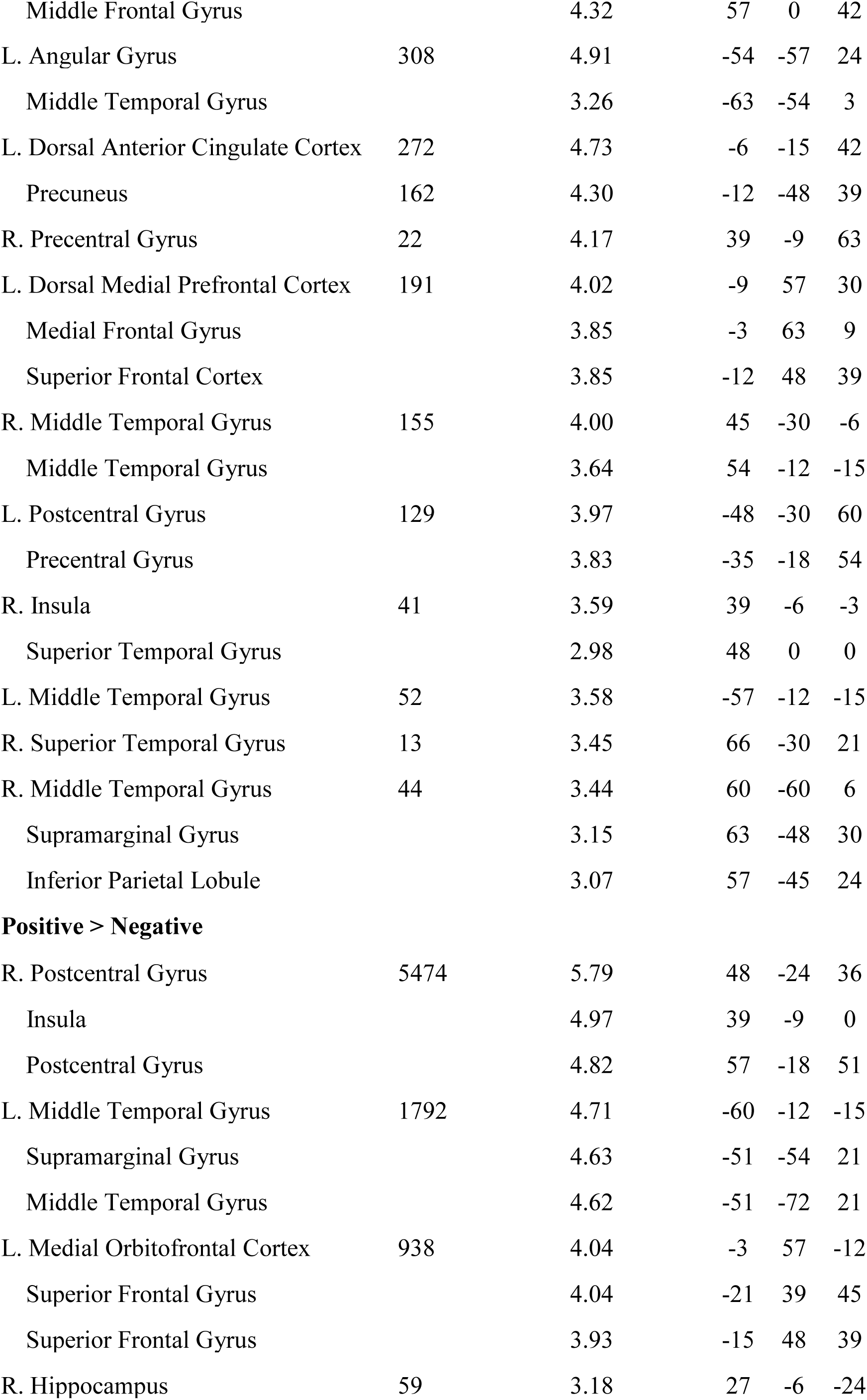
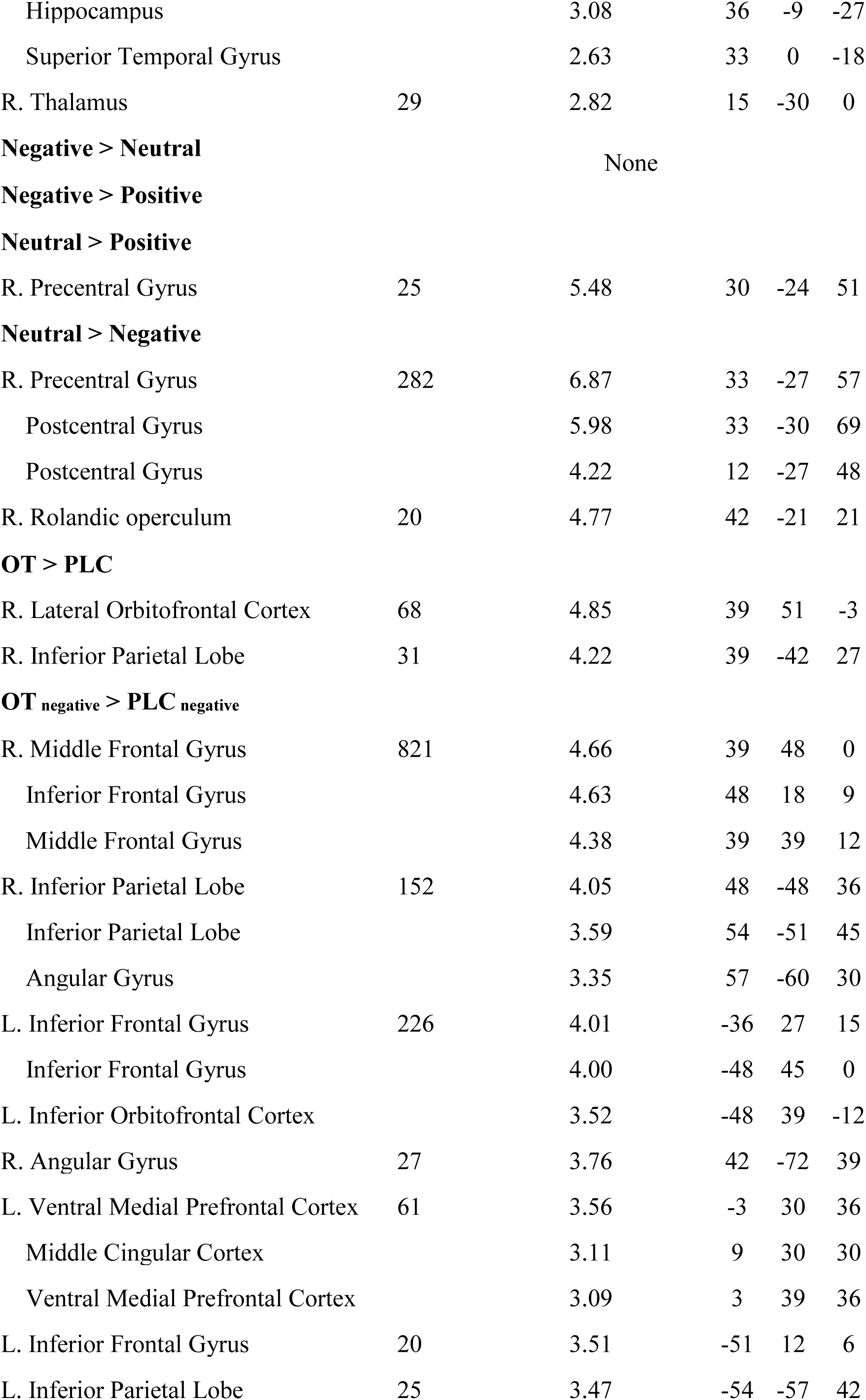
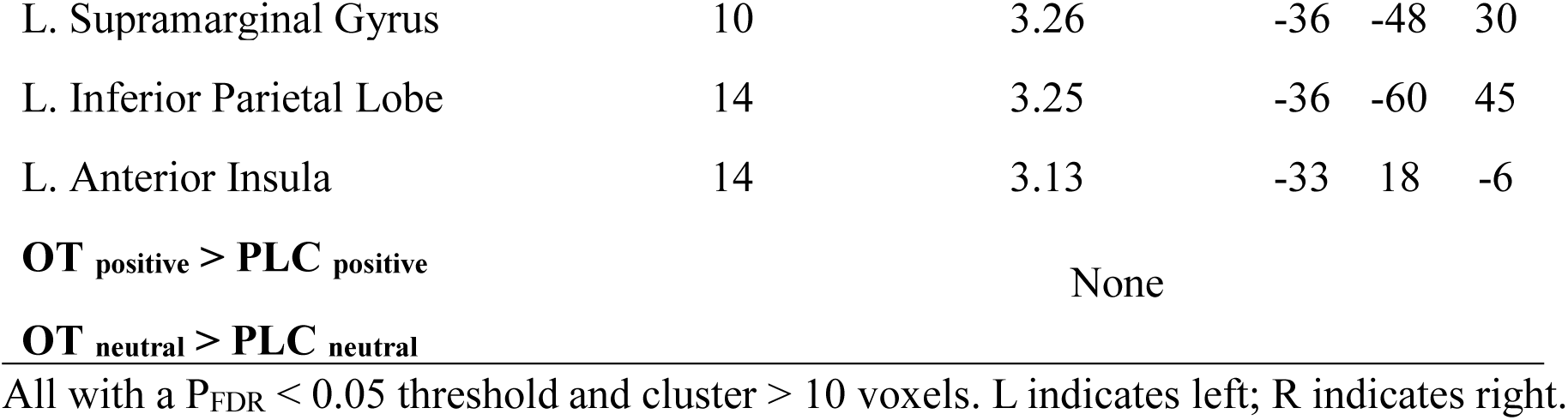
Significant activations induced by different touch stimuli at whole brain level.

**Figure 3.**
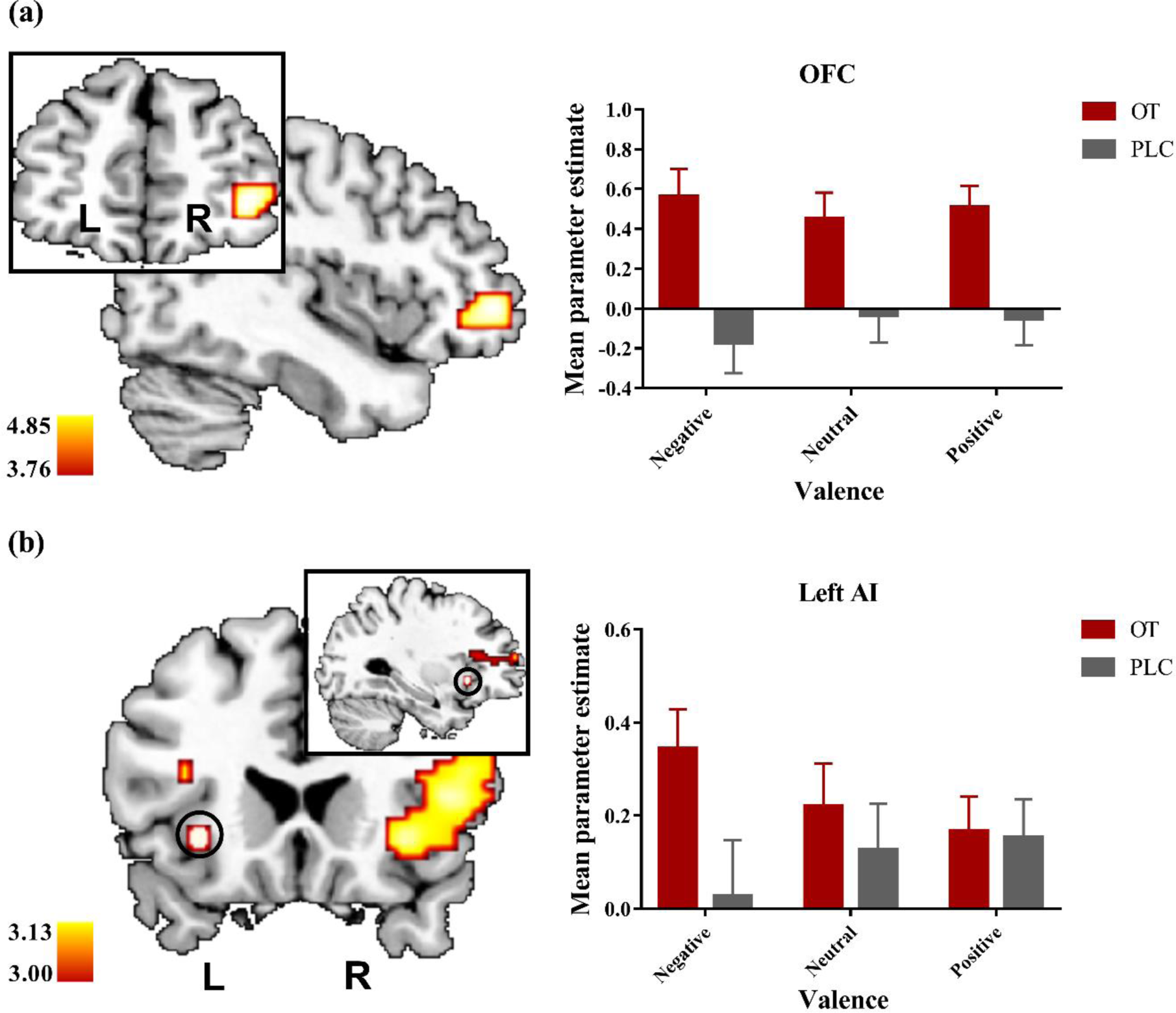
(a) The increased OFC (39 / 51 / −3) activity in response to affective touch across touch valences after the OT compared to PLC treatment during the whole brain analysis. (b) OT specifically enhanced neural responses to negative touch in the left AI (−33 / 18 / −6) compared to PLC treatment. Statistical maps are displayed with a height threshold of P < 0.001 uncorrected. Error bars show standard errors.

A functional connectivity analysis was performed using a generalized psychophysiological interactions (PPI) approach (gPPI; https://www.nitrc.org/projects/gppi; McLaren et al, 2012). Firstly, based on whole brain results, seed regions were defined as 6-mm spheres centered at the maximally activated voxel within the OFC and AI. One-sample t tests were used to examine OT’s modulatory effects on connectivity patterns during touch. No connections could survive the correction threshold of P_FWE_ < 0.05. Secondly, treatment effects involving the predefined ROIs were analyzed. Results revealed an increased functional connectivity between bilateral lateral amygdala and right AI in response to the negative relative to neutral touch (OT _neg > neu_ > PLC _neg > neu_, t = 4.58, P_FWE_ = 0.037, located in MNI space at 36 / 18 / −12, cluster size = 71). Extracted parameter estimates are given in Figure 4.

**Figure 4.**
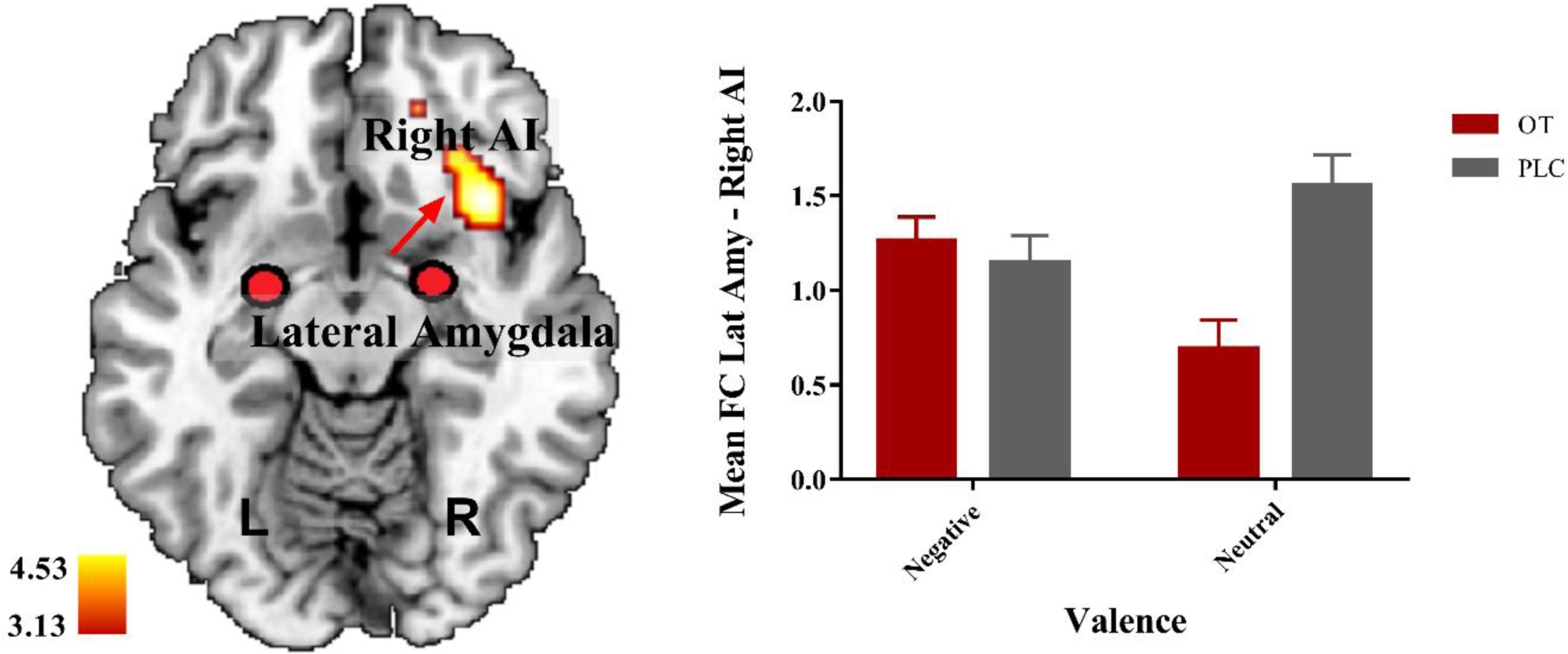
Increased bilateral lateral Amygdala – right AI functional connectivity to negative relative to neutral touch after the OT compared to PLC treatment. Statistical maps are displayed with a height threshold of P < 0.001 uncorrected. Error bars show standard errors.

### 3.4 Associations between basal OT concentrations and trait autism and on the effects of intranasal OT on responses to affective touch

Associations between basal OT concentrations and trait autism (AQ and STQ) and behavioral and neural responses to affective touch were calculated using Spearman rank correlation. Resuts showed that AQ scores were significantly negatively associated with baseline concentrations of OT (r = −0.314, P = 0.048). For behavioral and neural effects of OT we analyzed correlations between difference scores (i.e. OT-PLC) and plasma OT concentrations focussing only on significant effects of OT. There was a significant positive correlation between mean OFC responses across touch valences and basal OT concentrations (r = 0.417, P = 0.007) and also for anterior insula responses to negative valence touch (r = 0.347, P = 0.028) but not for pleasantness ratings across valences (r = 0.018, P = 0.911) or amygdala-AI functional conectivity in response to negative relative to neutral touch (r = – 0.152, P = 0.348) (see Figure 5). There were no significant correlations between AQ and STQ scores with OT-PLC difference scores for behavioral ratings and neural effects (Ps > 0.16).

**Figure 5.**
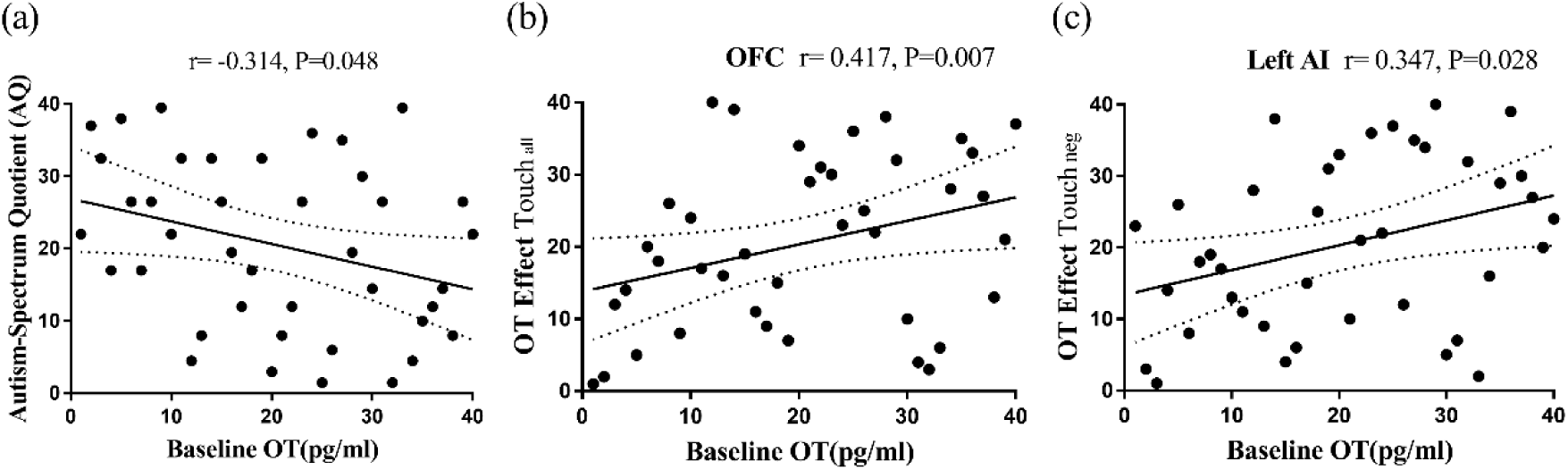
The basal concentrations of OT were (a) significantly negatively correlated with AQ scores and positively associated with neural effects of OT on (b) mean OFC responses across touch valences and (c) anterior insula responses to negative valence touch. The dotted line represents the 95% confidence interval.

## 4. Discussion

In the current study we investigated whether OT modulates behavioral and neural responses to affective touch administered with different valences and with no social intention. As hypothesized, intranasal OT significantly increased subjective pleasantness ratings of touch independent of valence and this behavioral effect was accompanied by enhanced neural effects on OFC responses across touch valences at the whole brain level. Furthermore, the OT-induced enhancement of OFC responses was positively associated with basal concentrations of OT in blood. Analyses on the effects of different individual touch valences further demonstrated that OT specifically increased AI responses to negative valence touch at the whole brain level and increased functional connectivity between the AI and amygdala for negative versus neutral valence. The increased AI responses were also positively correlated with basal plasma concentrations of OT. An exploratory analysis of behavioral pleasantness ratings also revealed that OT particularly increased them in the negative valence condition.

Analyses of the behavioral ratings of different touch valences revealed that, as expected, subjects perceived the positive touch as the most pleasant followed by the neutral and negative touch conditions. This finding therefore validated the effectiveness of the affective touch manipulation paradigm. At the neural level, consistent with previous findings in both healthy (Björnsdotter et al., 2014, 2010; Morrison, 2016; Olausson et al., 2016) and autistic patient (Croy et al., 2016; Kaiser et al., 2016) populations, positive valence touch induced stronger activation in the mPFC, IFG, STS, insula and ACC compared to neutral touch. Furthermore, enhanced activity in a region of the precentral gyrus induced by the neutral touch compared with positive and negative touch conditions also replicated previous findings using similar touch stimuli (Rolls, 2010; Rolls et al., 2003).

For the modulatory effect of OT relative to PLC on affective touch, analyses at the behavioral level showed that it enhanced pleasantness ratings across all touch valences and this was paralleled at the neural level by increased OFC activation, and that the latter were positively associated with prior basal OT concentrations in blood. Increased OFC activity following OT treatment has also been reported in a previous study in which positive interpersonal social touch was used (Scheele et al., 2014). Our current results at both the behavioral and neural level suggest that OT can enhance OFC responses to affective touch in a context where touch is administered with no social intention and across different valences. This finding is also in line with previous reports that both positive pleasant massage (Li et al., 2019) and painful/aversive responses (Boll et al., 2018) to tactile stimuli can increase endogenous OT release. Furthermore, OT can have analgesic effects and reduce the perceived unpleasantness of stimuli (see Boll et al., 2018; Walker et al., 2017). Given the role of OFC in encoding hedonic pleasure (Lamm et al., 2015; Mcglone et al., 2012; Rolls et al., 2003) and the increased pleasantness experienced by the subjects, OT may exert similar effects on affective touch by either potentiating the rewarding value of positive touch or reducing the perceived aversion of negative touch. This conclusion is further supported by increased scores in the positive mood and the decreased scores in the self-reported negative mood (supplemental information) across the affective touch paradigm under OT. Taken together, these findings demonstrate for the first time that OT has a general effect on enhancing the perceived hedonic value of non-socially intended affective touch independent of touch valence. Scheele et al. (2014) reported that OT increased pleasantness ratings when male subjects thought social touch was administered by a female rather than a male although the touch was actually delivered by the same female experimenter in all cases. Indeed, pleasantness ratings were much lower under PLC treatment in response to male compared with female touch indicating potential aversion to the male touch. Thus, valence may modulate the influence of OT when touch is socially administered whereas when there is no social intention such it may play a reduced role.

We confirmed previous findings of a negative association between autistic traits and basal OT concentrations in blood (Li et al., 2019; Zhang et al., 2016). Additionally, we observed that the effects of OT on increasing OFC activation to all touch valences were positively associated with basal blood OT concentrations. There was also a similar positive association between basal OT concentrations and the effect of intranasal OT on the anterior insula response to negative valence touch. This is the first evidence from healthy subjects for a potential modulatory influence of basal OT on responses in both brain reward and salience systems evoked by intranasal OT, although there is some previous evidence from clinical autism studies (Gordon et al., 2013a; Parker et al., 2017). Although it is not clear the extent to which basal OT concentrations in blood reflect those within the brain (Valstad et al., 2017), higher basal OT may increase sensitivity to exogenous administration by increasing receptor sensitivity. Indeed, lower basal OT concentrations in blood tend to be associated with lower OT receptor mRNA expression (Oztan et al., 2018).

Further disentangling of OT’s effects on each touch condition separately showed that at the neural level OT increased the AI response to negative valence stimuli at the whole brain level and an ROI-based analysis also revealed increased amygdala-AI functional connectivity for negative versus neutral valence. There was also some evidence from an exploratory analysis that at the behavioral level OT produced stronger effects on increasing pleasantness ratings for the negative valence touch. As a core hub of the salience network (Menon, 2015; Uddin, 2015; Uddin et al., 2017) and emotion processing (Koban and Pourtois, 2014; Kurth et al., 2010; Lamm and Singer, 2010), the AI has consistently been found to be involved in the modulatory effects of OT on social salience processing in various social tasks (Riem et al., 2011; Striepens et al., 2012; Yao et al., 2018a, 2018b). The increased strength of functional connectivity between the lateral amygdala and right AI also suggests an effect of OT on the salience network for negative relative to neutral valence touch. Indeed, OT has previously been shown to differentially influence functional connectivity between the AI and amygdala dependent upon both sex and salience (Gao et al., 2016). From this perspective, OT may have particularly increased the salience of the negative touch stimulus in view of its effects on the OFC altering it from being experienced as aversive, and therefore to be avoided, to becoming mildly pleasant and acceptable. The insula and its functional connectivity with the amygdala may be particularly important for encoding such valence shifts and indeed a number of brain regions have been shown to exhibit such an “affective mode” type of processing (see Berridge, 2019). On the other hand, the AI has also been shown to be important for prediction error encoding (Yao et al., 2018b) and a recent study has reported increased AI responses in response to tactile mismatch responses (Allen et al., 2016). Thus OT may have acted to increase a mismatch between an expected aversive response to the negative valence touch when the actual outcome was a mildly pleasant experience.

We were unable to demonstrate any effects of OT on responses to affective touch in other key regions involved in the processing of affective touch such as the STS and amygdala even using an ROI-based analysis. This may reflect the fact that we used non-socially directed touch applied via objects and indeed a recent study has reported that, in contrast to the AI, neutral object-based touch failed to activate these two regions compared to touch administered by others or by self (Boehme et al., 2019). Similar findings have been reported for the STS for hand compared to machine administered massage (Li et al., 2019).

The current study still has following limitations. Firstly, although one previous study has indicated both social touch and non-socially administrated tactile touch manipulated with the same velocity can induce similar perceived pleasantness and neural responses (Voos et al., 2013), the affective tactile touch stimuli we used may have relatively lower ecological validity than social touch. Secondly, the negative valence touch stimulus used did not evoke strong aversion and possibly OT would not have the same effect under such circumstances. Thirdly, only male subjects were included in the present study to avoid possible menstrual cycle effects. A number of recent studies have reported sex-dependent effects of OT in other domains (see Gao et al., 2016; Lieberz et al., 2019) thus whether the current findings would also occur in females is unclear.

In conclusion, the present study suggests that OT can facilitate responses to non-socially directed affective touch targeting CT fibers at both behavioral and neural levels and this effect is independent of touch valence. OT may increase the rewarding value of affective touch by influencing the OFC. Although OT exerted similar effects on different touch valences in the OFC, it additionally modulated responses to negative valence touch in the AI as well as its functional connectivity with the amygdala implying an additional effect on salience processing. These findings together provide evidence that OT may generally influence behavioral and neural responses to all types of touch which stimulate C-fibers, irrespective of valence and whether there is a social intention. Thus potentially OT may not only promote increased affiliation and approach behaviors following positive social touch but also more generally facilitate tactile exploration of the environment due to an increased liking for any kind of tactile stimulation.

## 5. Acknowledgments

This study was supported by the National Natural Science Foundation of China grant (NSFC grant number 31530032, 31700998). We thank Jianfu Li for the technical support.

