## supplemental materials for "Oxytocin increases the pleasantness of affective touch and orbitofrontal cortex activity independent of valence"

### Results

*Results of Personality and Mood Questionnaires*

The descriptive statistics on questionnaires showed that individual self-reported scores on anxiety, depression and social anxiety were within the normal healthy range (Table S1). To explore whether intranasal treatment would have effects on personal mood, we analyzed individual self-reported mood scores measured by the PANAS. A two way ANOVA analysis on PANAS scores was performed with treatment (OT vs. PLC) and time (time 1: pre-treatment; time 2: 30 min post-treatment; time 3: post-scan) as within-subject factors. For the negative mood scores, results showed a significant main effect of measured time ($F(2, 78) = 6.48$, $P = 0.007$, $\eta^2 = 0.14$), with a significant decrease of the negative mood after the scanning ($P = 0.02$, Cohen's $d = -0.45$) compared with the pre-treatment ratings scores. The main effect of treatment and the interaction effect were not significant ($Ps > 0.78$). Exploratory analysis found OT treated subjects rated lower negative mood scores after the scanning compared with pre-treatment ($P = 0.02$, Cohen's $d = -0.44$), but not in subjects with PLC ($Ps > 0.10$). For the positive mood scores, results revealed a marginally significant main effect of time ($F(2, 78) = 3.08$, $P = 0.06$, $\eta^2 = 0.07$). There were no significant main effect of treatment and interactions ($Ps > 0.57$). Exploratory analysis showed the positive mood scores significantly increased after the scanning relative to post-treatment rating scores in OT administrated subjects ($P = 0.04$, Cohen's $d = 0.32$), but not for the PLC group ($Ps > 0.99$; Figure S1).

### Supplemental figures

**Table S1.** Descriptive statistics of questionnaire scores (mean ± sd).

| Measurements | Rang | Mean ± sd |
|---|---|---|
| Beck Depression Inventory II | 0 - 25 | 7.74 ± 7.44 |
| State-Trait Anxiety Inventory | | |
| - State anxiety | 25 - 64 | 37.59 ± 8.55 |
| - Trait anxiety | 27 - 61 | 41.28 ± 8.16 |
| Liebowitz Social Anxiety Scale | | |
| - Fear | 4 - 48 | 21.03 ± 11.08 |
| - Avoid | 6 - 40 | 18.23 ± 9.32 |
| Empathy Quotient | 15 - 69 | 38.15 ± 10.72 |
| Autism Spectrum Quotient | 10 - 33 | 20.35 ± 5.58 |
| Social Touch Questionnaire | 18 - 60 | 39.98 ± 7.22 |

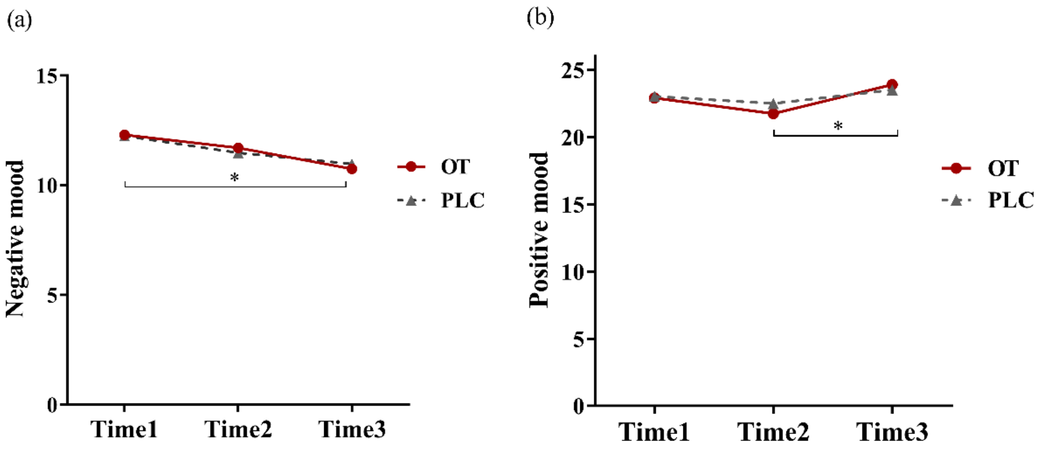


**Figure S1.** Individual mood scores of (a) the negative and (b) positive affection scale before and after the administration as well as after the scanning. $^{*}P < 0.05$.
